# The silence of the rewards: Non-reward cue suppresses cue-driven behaviour in pavlovian-instrumental transfer

**DOI:** 10.64898/2026.02.12.705619

**Authors:** Felippe E. Amorim, Sharon Morein-Zamir, Amy L. Milton

## Abstract

Flexible reward-seeking relies upon an organism’s integration of expectations about environmental cues with knowledge of actions that produce rewards, which is captured by pavlovian instrumental transfer (PIT). Although widely used to study motivation across species, rodent PIT paradigms differ from human tasks in lacking an active baseline for comparison. Here we adapted the full PIT paradigm to include a non-rewarded cue (CSØ) as an active control with the aim of improving the translational relevance of the procedure. Despite robust pavlovian and instrumental learning, neither male nor female rats expressed outcome-specific PIT when the active baseline was present. The absence of PIT persisted even after instrumental extinction training prior to the test, though cue-driven magazine approaches remained elevated. Data obtained under these conditions may reflect protocol-dependent modulation by the amount of training, or strain-specific differences in motivational influence. Replicated across male and female cohorts, they identify boundary conditions where PIT appears less robust. This methodological adaptation advances the translational alignment between animal and human PIT paradigms and informs future investigations into the mechanisms of cue-motivated behaviour.

## Introduction

In daily life, humans and animals integrate information from environmental cues dynamically to help their decision-making in obtaining rewards or avoiding punishments. For example, after a positive experience in a restaurant, appetitive cues subsequently such as seeing the outside of this establishment, indirectly influence behaviour by triggering a desire for the food (i.e. appetitive outcome) directly priming an action to go there (i.e. instrumental response). These everyday processes reflect fundamental and dynamic learning mechanisms, and how they occur is a key question in psychology. To better understand the underlying learning mechanisms, psychologists often use paradigms that isolate stimulus, outcomes and instrumental response associations (Balleine and Dickinson, 1998). Among the theoretical frameworks available, pavlovian instrumental transfer (PIT) is a central paradigm to understand how environmental reward-paired cues influence behaviour in both humans and animals.

Within PIT, two distinct psychological phenomena can be studied: general and specific. General PIT enhances animals’ behavioural actions associated to the same motivational domain when a predictive cue is given, whereas specific PIT helps them choose actions leading to the exact predicted outcome (specific-same), and discouraging those associated with different outcomes (specific-different). Continuing the restaurant example, cues associated with the establishment may make a person feel hungry and inclined to seek food (general PIT). At the same time, these cues may selectively promote actions leading to the previously rewarding dish (specific-same), while reducing the motivation for other menu items associated with different past experiences (specific-different). Animal studies evaluating these three transfer effects simultaneously – the ‘full transfer paradigm’ – have provided insights into the neural substrates associated with each type of transfer effect (Corbit and Balleine, 2005). Although this paradigm has some similarities to the ones used during human experimental approaches, it lacks the presentation of a non-rewarded paired cue used as a baseline in humans (Quail et al., 2017), posing fundamental challenges to cross-species translation.

In PIT paradigms, pavlovian and instrumental associations with an appetitive outcome are trained in separate sessions, followed by a test measuring the instrumental response in the presence of these pavlovian cues (**Fig. 1**). Most animal studies use one of two approaches to study PIT; either a single-lever or an outcome-specific paradigm (Cartoni et al., 2016). For the first, the presentation of a paired pavlovian cue enhances instrumental responses, in comparison with an unpaired conditioned stimulus (CS) or neutral non-CS, with the latter used to evaluate only the instrumental response enhancement due to the reinforced CS (Holmes et al., 2010). It has been suggested that the single-lever paradigm evokes a general transfer which is linked to a habitual instrumental response, controlled by stimulus-response associations (Cartoni et al., 2013). On the other hand, the outcome-specific paradigm evokes the expression of specific transfer by pairing two CSs with different outcomes, and two levers with each of these outcomes. Corbit and Balleine (2005) developed the full transfer paradigm to evaluate simultaneously both general PIT and selective cuing properties of excitatory stimuli. Due to this paradigm, studies have successfully identified differential neural substrates associated with each transfer effect (Corbit and Balleine, 2005) and demonstrated that general PIT is susceptible to devaluation (Corbit et al 2011).

**Figure 1.**
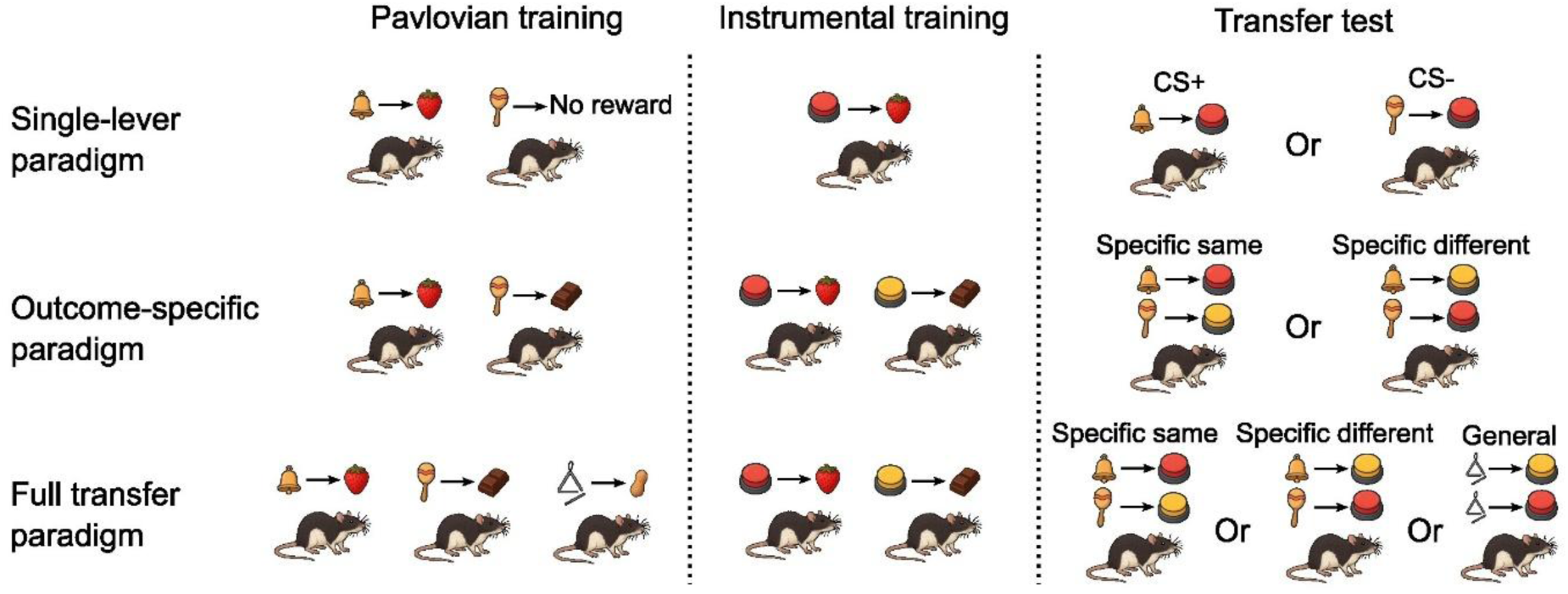
Schematic diagram of Pavlovian Instrumental Transfer (PIT) paradigms. Top row: In the single-lever paradigm, pavlovian training pairs one conditional stimulus (CS) with a reward and another with no reward. During the instrumental training, a single lever is associated with the outcome. In the transfer test, lever pressing is compared when presented with the reward-predictive (CS+) or non-rewarded cue (CS-). Middle row: In the outcome-specific paradigm, two pavlovian cues are each paired with distinct outcomes. Instrumental training involves two levers, each paired with one of the same rewards. During transfer, responding is assessed when CS and lever are paired with the same outcome (outcome-specific same) or when they are linked to different outcomes (outcome-specific different). Bottom row: In the full transfer paradigm, three pavlovian cues are paired with three distinct outcomes. Instrumental training involves only two of the rewards previously presented during the pavlovian phase, each paired to one lever. Lastly, the transfer test allows the measurement of outcome-specific effects and a general transfer effect, a non-specific effect that occurs when the pavlovian cue, which outcome was not paired with a lever, enhances the instrumental response.

Despite substantial progress made in understanding PIT, studies focusing on humans are relatively recent (Paredes-Olay et al., 2002, Hogarth et al., 2007, Talmi et al., 2008) in comparison with rodents (Estes, 1943, Estes, 1948, Lovibond, 1981, Corbit and Balleine 2003). Methodological and biological factors may therefore contribute to substantial differences in the evaluation of PIT in humans, compared with rodents. For example, gender can modulate the intensity of instrumental actions during the transfer phase in both appetitive and aversive contexts (Degni et al., 2024). Animal research using the full transfer paradigm lacks a non-paired reward cue, employed in human studies, to fully evaluate the facilitation of instrumental response (Quail et al., 2017). Without this evaluation, any instrumental performance change during the test phase could be due to inhibition during the baseline period, rather than an increase in motivation during CS presentation, or greater inhibition of incompatible actions rather than promotion of compatible actions (Quail et al 2017). Thus, there is a need to develop paradigms that better correspond to recent human designs.

In humans, most PIT studies have been conducted in predominantly female samples, and accumulating evidence suggests sex-differences in learning styles, social preferences, risk aversion and competitive behaviour (Severiens and Ten Dam, 1994; Falk and Hermle, 2018; Charness and Gneezy, 2012). Furthermore, the lack of analysis of sex differences in animal studies of cue-guided choice using the full transfer paradigm is also an issue that compromises the generalisation of the findings (Shansky and Murphy, 2021). More broadly, a lack of refined understanding of sex as a biological variable in behavioural strategies and underlying neural pathways complicates the translation of PIT findings into clinical applications. Studies using a specific transfer paradigm with two levers and two CSs reported mixed results with similarities in specific transfer effects (Alarcón and Delamater, 2019), but with female rats being more resistant to PIT extinction than males (Delamater et al., 2017). These findings suggest that cue-guided behaviour, at least for specific transfer, may be sex-dependent. Together, these design-related and sex-related factors hinder the translational validity of PIT findings.

In the present study, we devised a new strategy adapting the full transfer paradigm to improve the translation of the findings. In the original protocol proposed by Corbit and Balleine (2005), three CS-reward pairings are used with one of them used to measure general PIT effects. We first replaced it for a CS paired with no reward (CSØ), which enabled the comparison of PIT effects with a non-paired control CS. Using this approach, we hypothesized that during CSØ presentation, like studies with humans (Quail et al., 2017), rats would have a similar performance to the baseline. We further measured sex differences using female Lister-Hooded rats in this cue-guided behavioural task during the pavlovian and instrumental training phases.

## Materials and methods

### Subjects and apparatus

Subjects were 24 male Lister-Hooded rats for Experiment 1 (weighing 288-377g at the start of experiments), 12 Male Lister-hooded rats for Experiment 2 (weighing 305-375g at the start of experiments) and 24 female Lister-Hooded rats for Experiment 3 (weighing 192-237g at the start of experiments; all Charles River, UK). Animals were housed together in groups of 3 rats of the same sex per cage with environmental enrichment, and acclimatised to the animal facility for at least 7 days prior to experiments. During this time access to food and water was *ad libitum*. Prior to the start of behavioural procedures, the animals were food-restricted such that they were maintained at 90-95% of their age-matched free-feeding weight, being fed after the completion of each day’s behavioural procedures. This was in addition to any food earned during the behavioural procedures. Furthermore, at least one handling session was conducted before experiments to habituate animals and reduce stress. Five female rats were subsequently excluded due to audiogenic seizures that occurred during training sessions. This research was regulated under the UK Animals (Scientific Procedures) Act 1986 Amendment Regulations (2012) on Project Licences PA9FBFA9F and PP9536688 following ethical review by the University of Cambridge Animal Welfare and Ethical Review Body.

Twelve operant conditioning chambers (Med Associates, St. Albans, VT, USA) were used, equipped with a clicker and two audio generators (2.9KHz tone and white noise) inside sound- and light-attenuating chambers. Each chamber was illuminated by a white houselight during the experimental sessions. Depending on the training session, a pellet dispenser delivered a sucrose or a chocolate-flavoured pellet (AIN-76A, Sandown Scientific, Middlesex, UK). Ten of each pellet was put inside the home cages to reduce taste neophobia, at least one day before the first day of the experiment.

### Behavioural procedures

The full transfer paradigm experimental design is outlined in **Table 1**. The behavioural procedure took eighteen days (5 consecutive days per week) with nine days for pavlovian training, followed by seven days of instrumental sessions and subsequently, two days of transfer test. The procedure was performed in a schedule of 5 days of training per week. For the second experiment, we included one day of instrumental extinction prior to the transfer test.

**Table 1.** Experimental design for the full transfer paradigm used in all experiments. CS: conditioned stimulus; O: outcome; R: Response.

|  |  |  |
| --- | --- | --- |
| CS1-O1 | R1-O1 | CS1: R1 vs R2 |
| CS2-O2 | R2-O2 | CS2: R1 vs R2 |
| CSØ-OØ |  | CSØ: R1 vs R2 |

#### Pavlovian training

Rats performed nine days of pavlovian training with two sessions per day, each session for a different reward. Sessions were separated by at least two hours. A combination of three different auditory stimuli (experiment 1 and 3: white noise, 1Hz clicker and tone, 74dB; counterbalanced across rats; experiment 2: 1Hz clicker, 10Hz clicker and tone, 74dB; counterbalanced across rats;) were used as conditioned stimuli (CS1, CS2 and CSØ) paired with two possible positive outcomes, sucrose or chocolate pellets (O1 or O2) or no pellets as a neutral outcome (OØ). The order of CS presentation was counterbalanced each day. During each session, a two-minute-long auditory stimulus was presented four times each in a random order, paired with its respective outcome (sugar or chocolate pellets, or no outcome for the CSØ). Each cycle consisted of an intertrial interval, CS and neutral stimulus, and within the session, each cycle started with a different component than the end of the previous cycle.

During the two-minute presentation, the relevant outcome was delivered using a random time 30-second (RT-30s) schedule of reinforcement. The intertrial interval was programmed to be the same as the stimulus presentation, two minutes.

#### Instrumental training

Rats performed two sessions per day, with one session presenting a single lever (R1) with one outcome (O1), and the other session presenting the other lever (R2) with the other outcome (O2).This was counterbalanced across animals, with the order of sessions (R1→O1 or R2→O2) alternated each day and with an interval of two hours between sessions. Each reward was delivered in a continuous reinforcement (fixed ratio 1, FR1) schedule for the first day. For days 2-4, the schedule was changed to a random-ratio-5 (RR-5) and for days 5-7, rewards were delivered on an RR-10 schedule.

For Experiment 3, on the day after the last instrumental training session, we performed two 30-minute instrumental extinction sessions, one for each lever. No appetitive reward was delivered with the lever press.

#### Pavlovian-instrumental transfer test

Rats performed two 20-minute transfer test sessions, one on each lever, on two separate days. Each session was non-reinforced, with one lever (R1 or R2) present throughout the whole session with no reward being delivered. A single lever was available during the first four minutes, before presenting any CS. The rest of the session consisted of delivering six two-minute trials (3 trials per CS), in which a CS paired with a positive reward (CS1 or CS2) or with a neutral stimulus (CSØ) could be presented. The second session of the day was similar, presenting the previously reinforced CS that had not been presented in the first session (counterbalanced).

Four different behaviours were calculated based on the lever response per trial. First, we measured the number of lever presses during the intertrial interval, which was a CS-free period between trials. For specific transfer, it was considered the “same” condition when the auditory CS promoted a lever response that shared the same outcome (i.e. CS1→R1 or CS2→R2). When the delivered CS predicted a different pellet, but the animal still performed a lever press (i.e. CS1→R2 or CS2→R1), this behaviour was considered a “different” outcome. The last category was ‘general transfer’ where the animal had an instrumental response to the auditory stimulus paired with no food (i.e. CSØ →R1 or CSØ→R2). All these behaviours were averaged per trial.

#### Data analysis

Data are presented as means ± standard error of the mean (S.E.M.). One-way repeated measures analysis of variance (ANOVA) followed by Bonferroni’s multiple comparison correction was applied to the instrumental training and transfer test data. For two-way repeated measures and three-way repeated measures and mixed ANOVAs, Mauchly’s test was applied to check the assumption of sphericity, with degrees of freedom corrected using the Greenhouse–Geisser estimates of sphericity in case this assumption was violated. Tukey post-hoc comparison was used for between subjects. A timescale of days (2 sessions per day) was considered for the pavlovian and instrumental phases because only one reward type was delivered during a single session. All statistical tests were two-sided with a significance level set at α = 0.05. The effect size was reported using a partial eta-squared value (η_p_2).

## Results

### A neutral non-conditioned stimulus inhibited outcome-specific effects

First, we evaluated performance during the pavlovian training phase by nosepoke entries and nosepoke duration (**Fig. 2A** and **2B**). Rats nosepoked more during the CSs than during the CSØ or ISI [Stimuli: (*F*_(1.21,27.8)_ = 131, *p* < .001, η_p_^2^ = .85], with responding for the CSs increasing across time [Session: *F*_(2.63, 60.4)_ = 11.0, *p* < .001, η_p_^2^ = .32; Stimuli x Session: *F*_(4.14, 95.2)_ = 14.2, *p* < .001, η_p_^2^ = .38]. Similarly, rats spent longer nosepoking during presentation of the reward-associated CSs [Stimuli: *F*_(1.54, 35.5)_ = 109, *p* < .001, η_p_^2^ = .83] with this time increasing for the CSs specifically as the sessions progressed [Session: *F*_(3.18, 73.0)_ = 7.96, *p* < .001, η_p_^2^ = .26; Stimuli x Session: *F*_(5.73, 132)_ = 8.28, *p* < .001, η_p_^2^ = 0.27]. Thus, rats learned that the CSs were predictive of reward across sessions, being more likely to enter the magazine when the CSs were presented, and spending longer in the magazine during these CSs.

**Figure 2.**
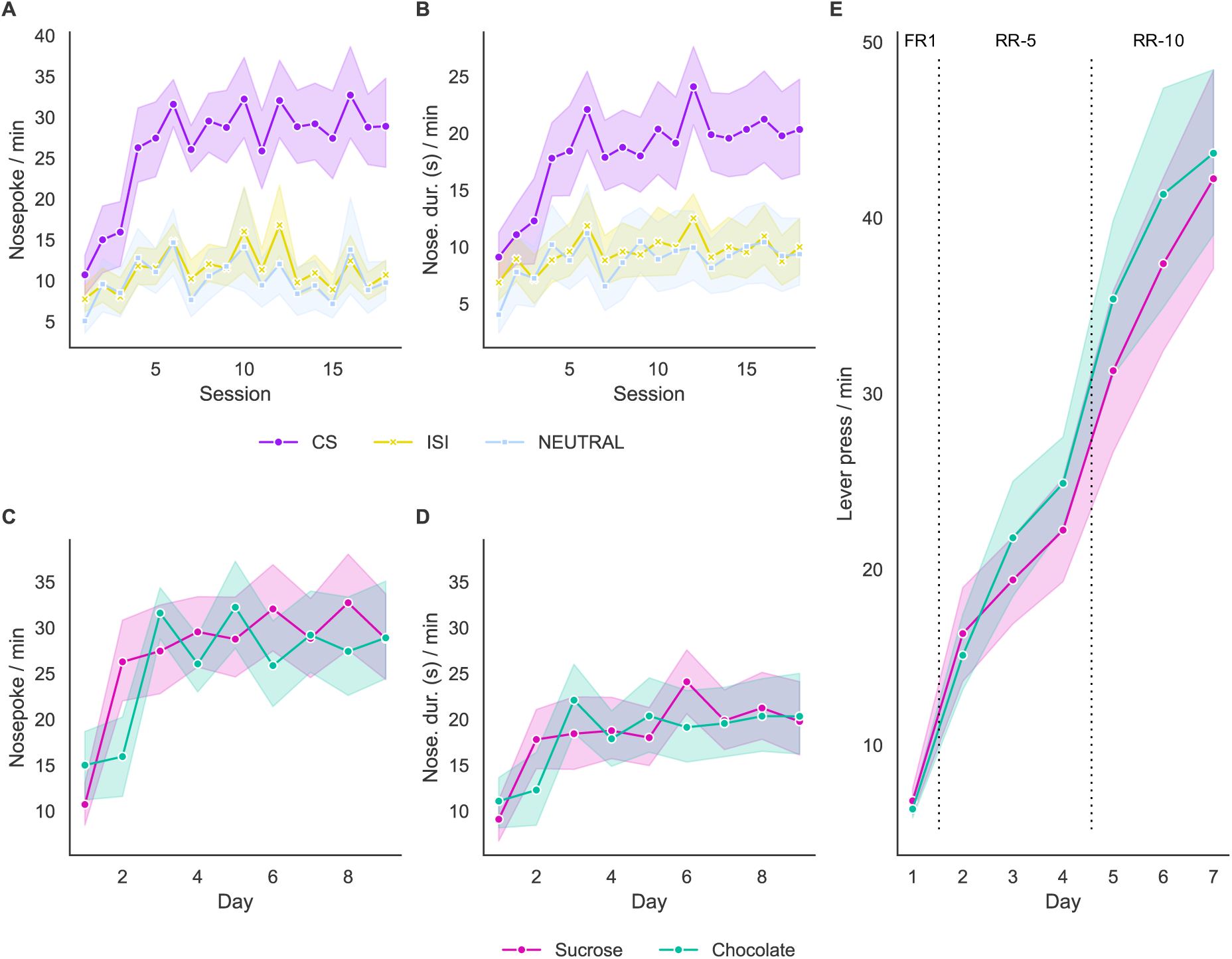
Performance during the pavlovian and instrumental phases across sessions during the first experiment. (A) Nosepoke entries and (B) duration according to the stimuli. CS group represents the performance when an auditory stimulus paired with a reward was delivered. ISI represents the interstimulus interval and NEUTRAL when a stimulus paired with no reward was delivered. (C) Nosepoke entries and (D) duration according to the reward offered. (E) Total amount of lever presses across seven days of the instrumental training. Sample size n = 24. Data are shown as mean ± 95% C.I.

As we had combined the CSs for the purposes of analysis of pavlovian conditioning, we further evaluated if animals exerted a reward preference that might have influenced nosepokes (**Fig. 2C**) or nosepoke duration (**Fig. 2D**) during the pavlovian conditioning. A two-way repeated measures ANOVA with factors of Time (days 1 to 9) and Reward (sucrose vs chocolate) showed that although rats increased their nosepoking across sessions [Time: *F*_(2.54, 58.3)_ = 20.8, *p* < .001, η_p_^2^ = .47] overall, nosepoking was equivalent for the two rewards [Rewards: *F*_(1, 23)_ = 1.27, *p* = .27]. However, there was a statistically significant interaction effect [Reward x Time: *F*_(4.81, 111)_ = 14.3, *p* < .001, η_p_^2^ = .38]. Pairwise comparisons for both rewards showed daily fluctuations in nosepoking, with no statistically significant differences for days 5 (*p* = .06), 7 (*p* = .83) and 9 (*p* = .96), while the sucrose condition had a higher performance for days 1 and 3, and chocolate condition for days 2, 4, 6 and 8. Note that in all experiments the order was counterbalanced across animals to mitigate any possible order effects.

Similarly, increases in nosepoke duration during both CSs were observed across sessions [Time: *F*_(2.51, 57.6)_ = 12.6, *p* < .001, η_p_^2^ = .35] with an overall similar preference for the two rewards [Reward: *F* < 1] though again a significant interaction [Reward x Time: *F*_(4.86, 111)_ = 8.33, *p* < .001, η_p_^2^ = .27]. Thus, although there were day-to-day fluctuations in reward preference, these were minor and the rats learned to discriminate between both reward-associated CSs, the CSØ and the ISI.

In addition, a two-way repeated measures ANOVA for the instrumental phase with Time and Reward as factors showed an increase in the lever presses across days (**Fig. 2E**; F(2.16, 49.56) = 143.24, p < 0.001, η2 =0.86), with no main effect of reward (F(1, 23) = 1.71, p = 0.20, η2 =0.07) and no time x reward interaction (F(3.52, 81.04) = 1.70, p = 0.17, η2 =0.07).

We assessed pavlovian-instrumental transfer in two separate tests, combined here for comparison (**Fig. 3**). Contrary to our predictions, rats did not change their responding during different cue periods (i.e., they responded no more during the reward-associated CSs than during the CSØ or the ITI) [**Fig. 3A**; Period: F_(2.62, 60.3)_ = 0.64, *p* = 0.59]. Bonferroni-corrected post-hoc tests revealed no significant difference between all PIT effects and control (p = 1 for ITI vs CSØ, same and different). To rule out any rapid extinction effects, we also analysed measured the mean lever press performance per trial using a three-way repeated measures ANOVA depending on the PIT outcome and reward (**Fig. 3B** and **3C**). For specific PIT and reward, lever pressing was highly influenced by the order within the day (F_(1.56, 36.0)_ = 49.3, *p* < 0.001, η_p_² = .68), with day also exhibiting a modest but significant effect (F_(1, 23)_ = 6.82, *p* = 0.016, η_p_² = .23). Moreover, PIT alone did not significantly influence behaviour (F_(1, 23)_ = 1.47, p = 0.24, η_p_² = .06), however, a significant Order x PIT interaction (F_(1.50, 34.61)_ = 3.96, *p* = 0.039, η_p_² = .15) indicated that order-dependent changes were modulated by which PIT was been evaluated. Finally, no effect of the reward was observed (F_(1, 23)_ = 0.02, *p* = 0.9), suggesting that the conditioned stimulus did not significantly influence instrumental response.

**Figure 3.**
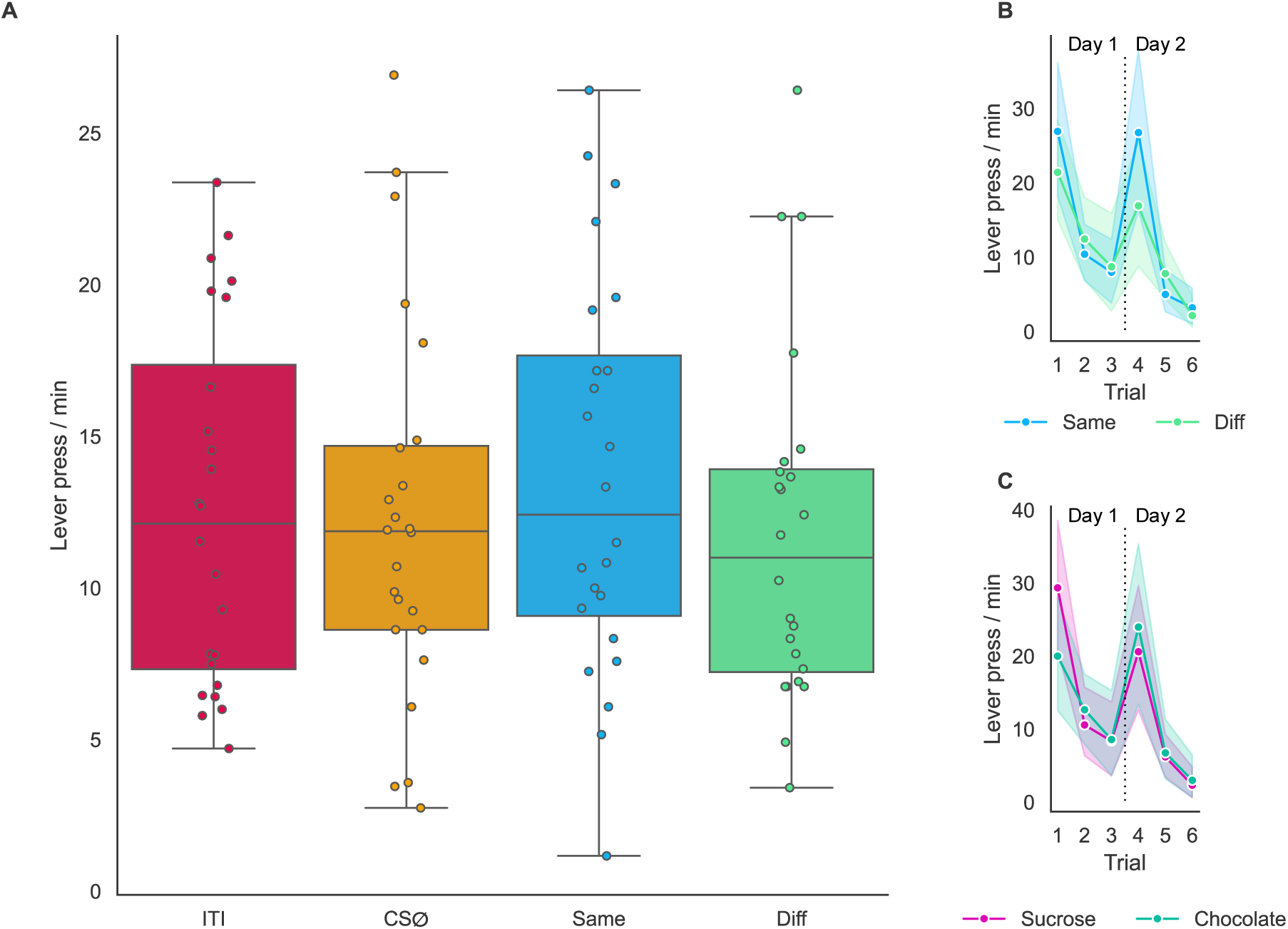
Animals showed neither general not specific transfer in a full transfer paradigm. Male Lister-hooded rats were trained on the full transfer paradigm PIT task and their performance was measured during the transfer phase. (A) The performance was classified based on the transfer effect, represented by intertrial interval (ITI), outcome-specific same, outcome-specific different (Diff) and general outcome. Lever presses total according to the PIT effect (B) and associated reward (C) across trials Sample size n = 24. Data are shown as mean ± 95% C.I.

Thus, despite good pavlovian and instrumental learning, rats showed evidence of neither specific pavlovian-instrumental transfer (i.e. no discrimination between the Same and Different outcomes at test) nor general pavlovian-instrumental transfer (i.e. an increase in responding to the CSØ). Moreover, rats did not increase responding above ITI levels, perhaps indicating that subtle pavlovian effects on responding were being masked by high levels of baseline instrumental behaviour.

### An extinction session prior to transfer test reduced test responding, but animals still did not show transfer effects

Although the training protocol was similar to that described in the literature, animals exhibited a higher total amount of lever presses per day during the instrumental training (last session: 2577.46±144.35 for both reinforcers) and across all transfer effects during test (ITI: 12.57±1.21; Same: 13.63±1.35; Different: 11.78±1.19; General: 12.29±1.29) in comparison with previous experiments from Corbit et al (2011). The apparent failure to replicate the specific PIT effects described by Corbit (2005, 2007) could be attributed to a ceiling effect in instrumental responding, which could be masking the enhancement effect of CS presentations. To reduce the amount of lever pressing during the transfer test and the potential influence of a ceiling effect, the procedure used in Experiments 2 and 3 included an extinction session one day before the transfer tests. Concomitantly, female rats on Experiment 3 were also used to assess sex differences during the learning process on the full transfer paradigm.

As for Experiment 1, male rats rapidly learned to nosepoke in the presence of reward-associated CSs [**Fig. 4A**; Stimuli: *F*_(1.09, 10.9)_ =59.6, *p* < .001, η_p_^2^ = .86], with nosepoking increasing selectively for the CSs across sessions [Time: *F*_(4.60, 46.0)_ = 4.40, *p* < .001, η_p_^2^ = .31; Stimuli x Time: *F*_(2.33, 23.3)_ = 7.88, *p* < .001, η_p_^2^ = .44]. Similar effects were observed for nosepoke duration, which was higher during the reward-associated CSs [**Fig. 4B**; Stimuli: *F*_(1.40, 14.1)_ = 104, *p* < .001, η_p_^2^ = .91] and increased selectively for the CSs across sessions [Time: *F*_(4.81, 48.1)_ = 1.26, *p* = .22; Stimuli x Time: *F*_(4.35, 43.5)_ = 2.36, *p* < .001, η_p_^2^ = .19].

**Figure 4.**
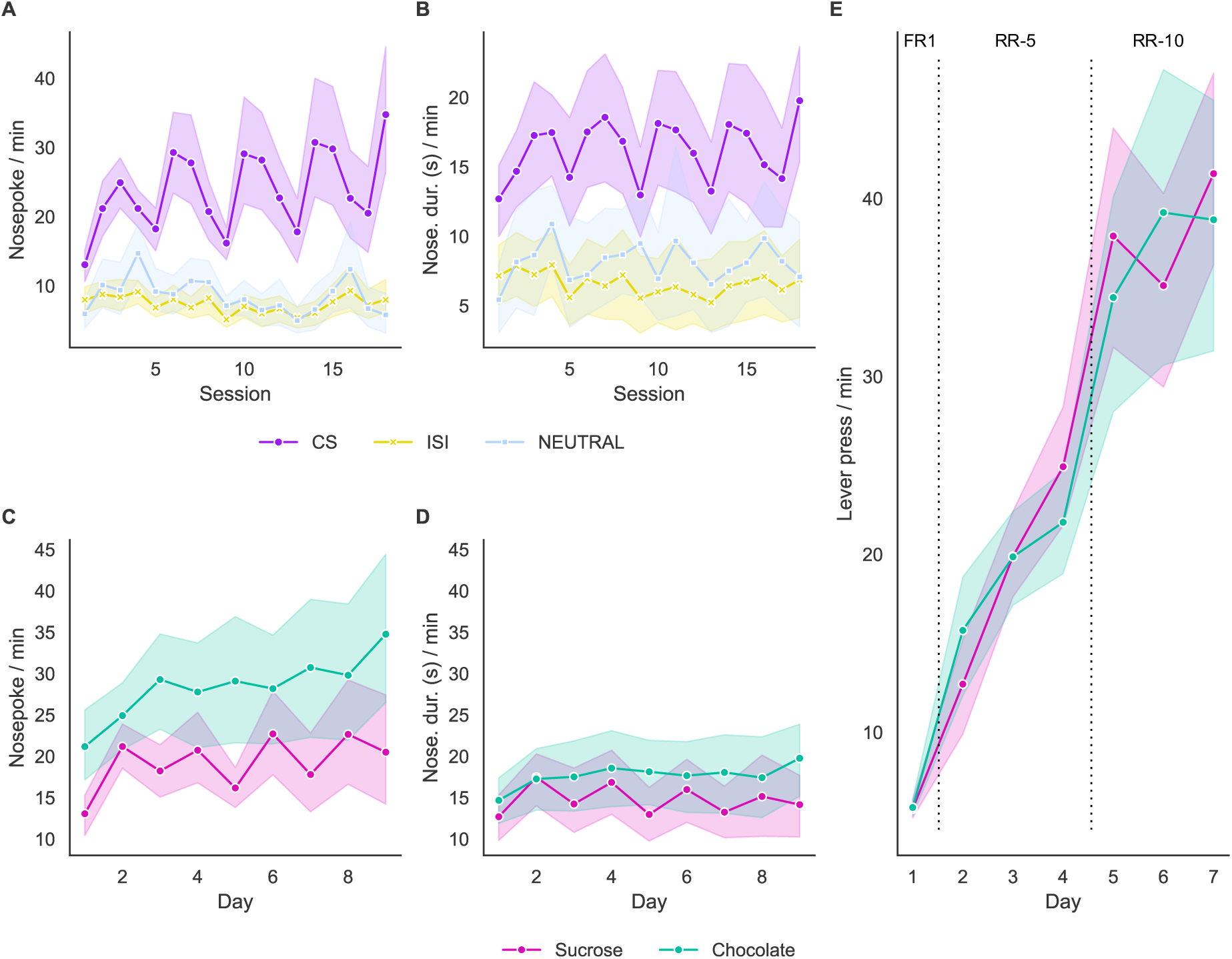
Performance of male rats for the pavlovian and instrumental phases across sessions during experiment 2. (A) Nosepoke entries and (B) duration according to the stimuli. (C) Nosepoke entries and (D) duration according to the reward offered. (E) The total amount of lever presses across seven sessions of the instrumental training. Sample size n = 12. Data are shown as mean ± 95% C.I.

To evaluate any reward preference, similar to Experiment 1, we measured the nosepoke entries (**Fig. 4C**) and duration (**Fig. 4D**) when an auditory stimulus associated with a reward was presented. Nosepoking increased across training as measured by magazine entries [Time: *F*_(2.86, 28.6)_ = 3.77, *p* = .001, η_p_^2^ = .27] and nosepoke duration [Time: *F*_(3.61, 36.1)_ = 1.52, *p* = .16, η_p_^2^ = .13]. However, this cohort showed a preference for chocolate over sucrose, which increased over the course of training. This was the case for both magazine entries [Reward: *F*_(1, 10)_ = 26.8, *p* < .001, η_p_^2^ = .73; Time x Reward: *F*_(3.60, 36)_ = 5.05, *p* < .001, η_p_^2^ = .34] and nosepoke duration [Reward: *F*_(1, 10)_ = 11.5, *p* = .007, η_p_^2^ = .18; Time x Reward: *F*_(3.92, 39.2)_ = 2.14, *p* = .04, η_p_^2^ = .18]. However, when working for reward (**Fig. 4E**), there appeared to be equivalent preference between chocolate and sucrose pellets, with instrumental responding increasing for both with training [Time: *F*_(2.51, 27.6)_ = 76.9, *p* < .001, η_p_^2^ = .87; Reward: *F* < 1; Time x Reward: *F*_(3.27, 36.0)_ = 1.26, *p* = .29].

To avoid the high levels of instrumental responding – and potential masking of PIT effects – seen in Experiment 1, we conducted an extinction session for both levers prior to the transfer test. Notably, animals reduced their lever pressing, which removes the possibility of a ceiling effect (**Fig. 5A**). However, as for the previous experiment, we did not observe an overall PIT effect (Period: *F* < 1). Rats did not modulate their responding based on whether they were receiving the Same or Different CS to the reward associated with the lever press [**Fig. 5B**; PIT: *F*_(1, 11)_ = 0.28, *p* = 0.61] though the order in which the CSs were presented across the two tests influenced responding [Order: *F*_(1.04, 11.4)_ = 11.69, *p* = .005, η_p_^2^ = .52; PIT x Order: F_(1.04, 11.4)_ = .17, *p* = .69]. Behaviour remained stable across days (Day: F_(1, 11)_ = .03, *p* = .86, η_p_² = .003). Moreover, response significantly depended on the reward presented (**Fig. 5C**; Reward: F_(1, 11)_ = 10.32, *p* = .008, η_p_² = .48). Notably, a significant interaction between reward, order and days (Reward x Order: F_(1.36, 14.93)_ = .99, *p* = 0.36, η_p_² = .082; Reward x Day: F_(1, 11)_ = 8.53, *p* = .014, η_p_² = .437; Reward x Order x Day: F_(1.13, 12.40)_ = 18.49, *p* < 0.001, η_p_² = .63), suggested a strong interactive effect that modulated instrumental response during the test for specific PIT.

**Figure 5.**
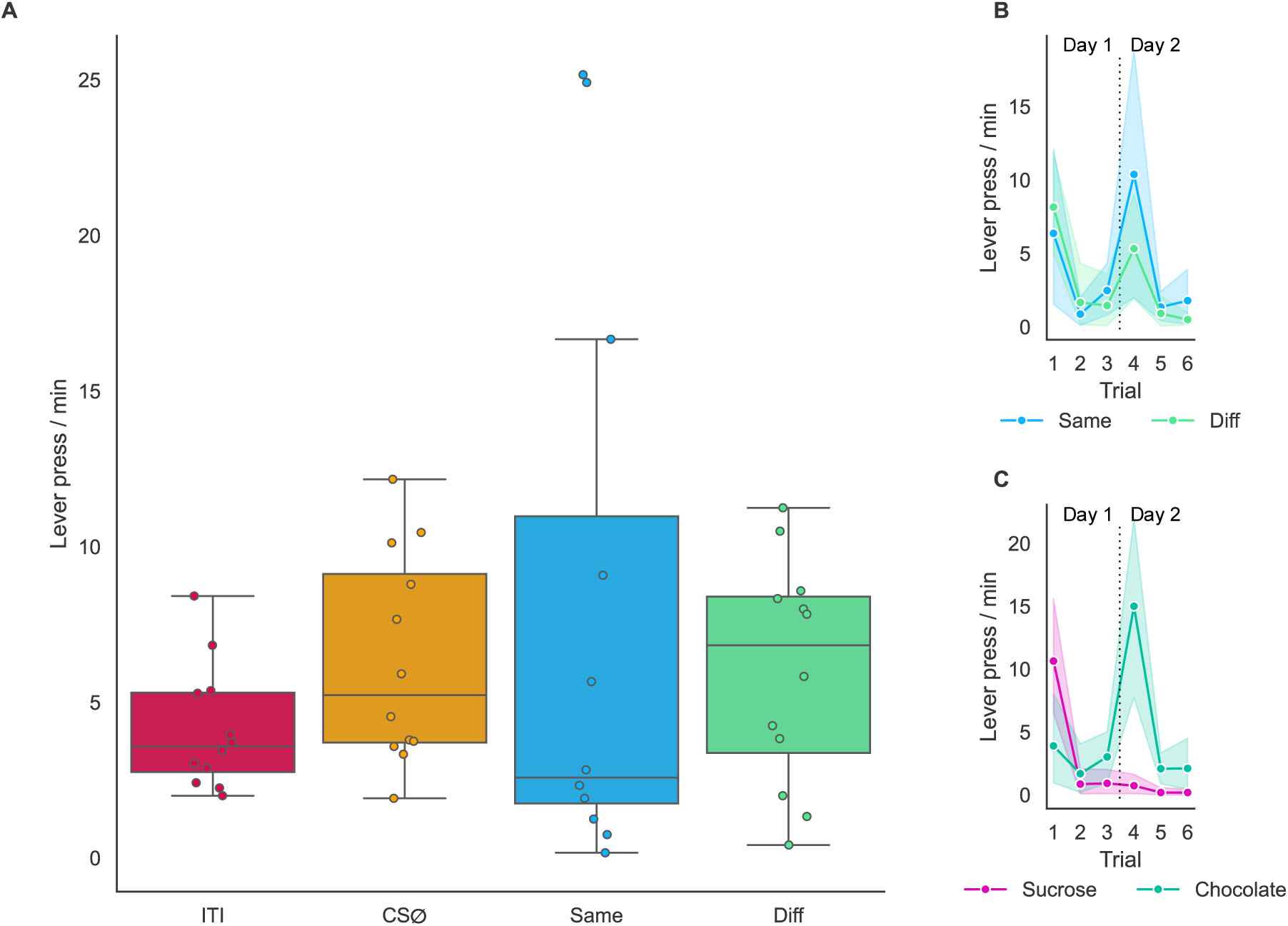
Instrumental extinction prior to the test reduces lever presses but male rodents did not show transfer effects in a full transfer paradigm. Male Lister-hooded rats were trained on the full transfer paradigm PIT task, performed one instrumental extinction session and their performance was measured during the transfer phase. (A) The performance was classified based on the outcome, represented by intertrial interval (ITI), outcome-specific same, outcome-specific different (Diff) and general transfer effect. Lever presses total according to the PIT effect (B) and associated reward (C) across trials. Sample size n = 12. Data are shown as mean ± 95% C.I.

Thus, as for Experiment 1, there was evidence of good pavlovian and instrumental learning that did not manifest as PIT effects during the transfer phase.

### Female Lister hooded rats learned the pavlovian and instrumental phases, but also did not show transfer effects

Using the same procedure as Experiment 2, it was also found that female rats showed consistent learning across the pavlovian training phase (**Fig. 6A** and **6B**). Rats increased the numbers of magazine entries selectively during the reward-associated CSs across the course of training [Time: *F*_(4.05, 72.8)_ = 10.8, *p* < .001, η_p_^2^ = .38; Stimuli: *F*_(1.13, 20.3)_ = 133, *p* < .001, η_p_^2^ = .88; Time x Stimuli: *F*_(5.78, 104)_ = 14.7, *p* < .001, η_p_^2^ = .45]. The duration of magazine entries also increased selectively for the reward-associated CSs [Time: *F*_(4.14, 74.6)_ = 4.55, *p* = .002, η_p_^2^ = .20; Stimuli: *F*_(1.29, 23.2)_ = 95.4, *p* < .001, η_p_^2^ = .84; Time x Stimuli: *F*_(5.49, 98.9)_ = 7.82, *p* < .001, η_p_^2^ = 0.30].

**Figure 6.**
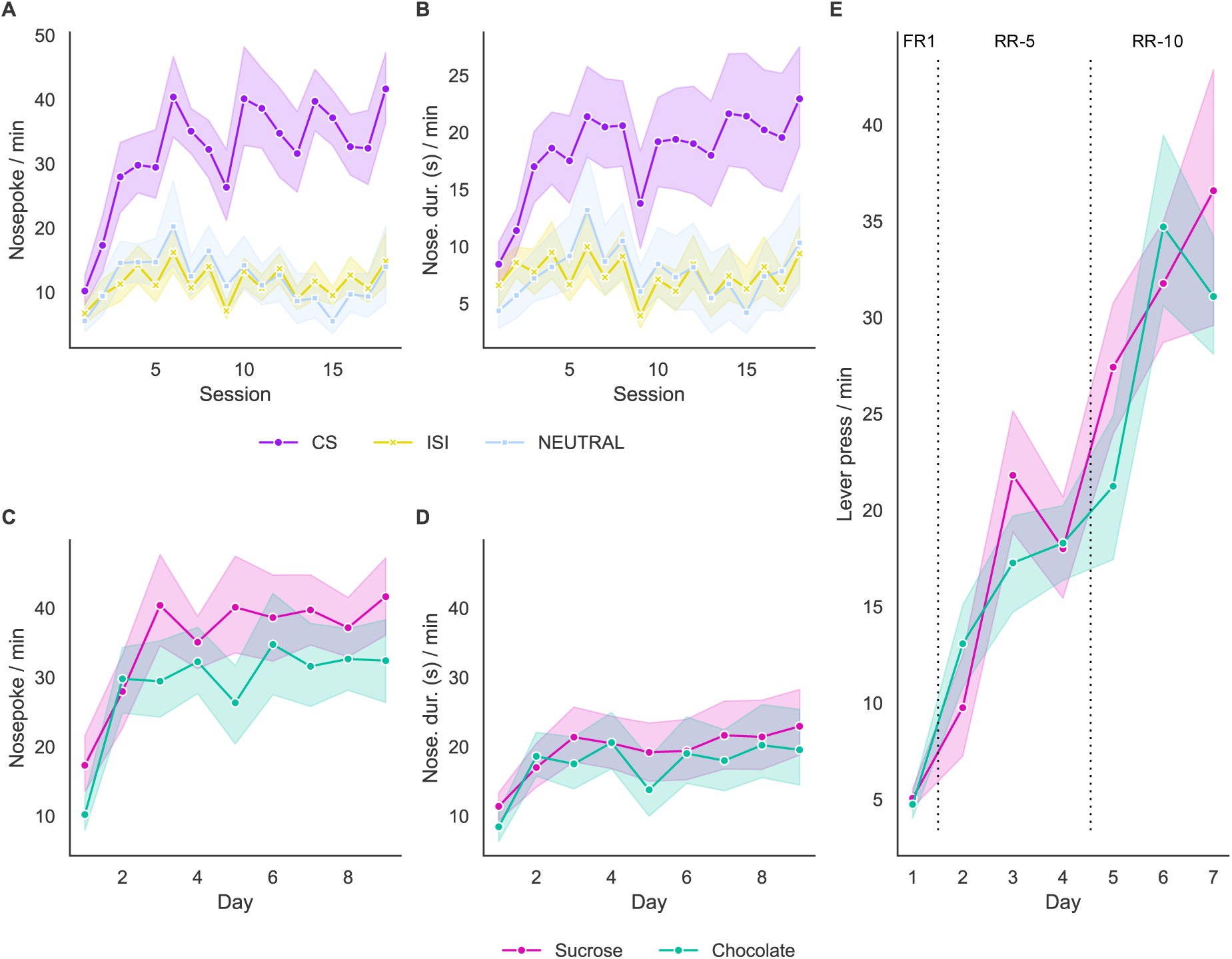
Performance for the pavlovian and instrumental phases across sessions during the second experiment. (A) Nosepoke entries and (B) duration according to the stimuli. (C) Nosepoke entries and (D) duration according to the reward offered. (E) The total amount of lever presses across seven days of the instrumental training. Sample size n = 19. Data are shown as mean ± 95% C.I.

Reward preference was also analysed for Experiment 3 (**Fig. 6C** and **6D**). The number of magazine entries made indicated that females had an overall preference for the sucrose over the chocolate pellets [Reward: *F*_(1, 18)_ = 22.1, *p* < .001, η_p_^2^ = .55], which increased over the course of training [Time: *F*_(3.59, 64.6)_ = 23.3, *p* < .001, η_p_^2^ = .56; Reward x Time: *F*_(4.26, 76.6)_ = 6.75, *p* < .001, η_p_^2^ = .27]. Nosepoke duration increased over the course of training [Time: *F*_(2.87, 51.7)_ = 9.72, *p* < .001, η_p_^2^ = .35] but slightly more for sucrose than chocolate [Reward x Time: *F*_(8, 144)_ = 4.03, *p* < .001, η_p_^2^ = .18]. Nevertheless, there was no overall difference in the time spent nosepoking for the two rewards [Reward: *F*_(1, 18)_ = 2.77, *p* = .11]. Finally, females learned the instrumental responses associated with the two rewards, increasing their responding across the course of training [**Fig. 6E**; Time: *F*_(2.82, 50.7)_ = 75.0, *p* < .001, η_p_^2^ = .80). Although there was no overall preference for either reward [Reward: *F*_(1, 18)_ = 1.83, *p* = .19], preference appeared to fluctuate across instrumental sessions [Time x Reward: *F*_(3.08, 55.4)_ = 6.10, *p* = .001, η_p_^2^ = .25]. These results indicate that the female rats learned both the pavlovian and instrumental associations during training, with the reward type having no significant effect on learning curves.

The rats underwent an instrumental extinction session prior to the transfer test. This led to lower levels of responding during the transfer test itself (**Fig. 7A**; ITI: 7.94±1.00; Same: 11.56±2.40; Different: 10.03±1.71; General: 12.00±1.68). However, as in the first and second experiments, there appeared to be no transfer effect, with similar levels of responding across all phases of the test [Period: *F*_(1.90, 34.2)_ = 1.03, *p* = 0.37]. Rats did not show differences in responding to the Same or Different stimuli (**Fig. 7B**; PIT: *F*_(1,18)_ = 0.18, *p* = .678, η_p_^2^ = .01). While the order of presentation influenced responding, this affected responding to the Same and Different specific PIT outcomes equivalently [Order: *F*_(1.47, 26.55)_ = 9.92, *p* = .002, η_p_^2^ = .36; PIT x Order: *F*_(1.49, 26.84)_ = .08, *p* = .867, η_p_^2^ = .005]. However, a significant PIT x Day interaction suggests that PIT varied across testing days [PIT x Day: *F*_(1, 18)_ = 6.75, *p* = .018, η_p_^2^ = .27].

**Figure 7.**
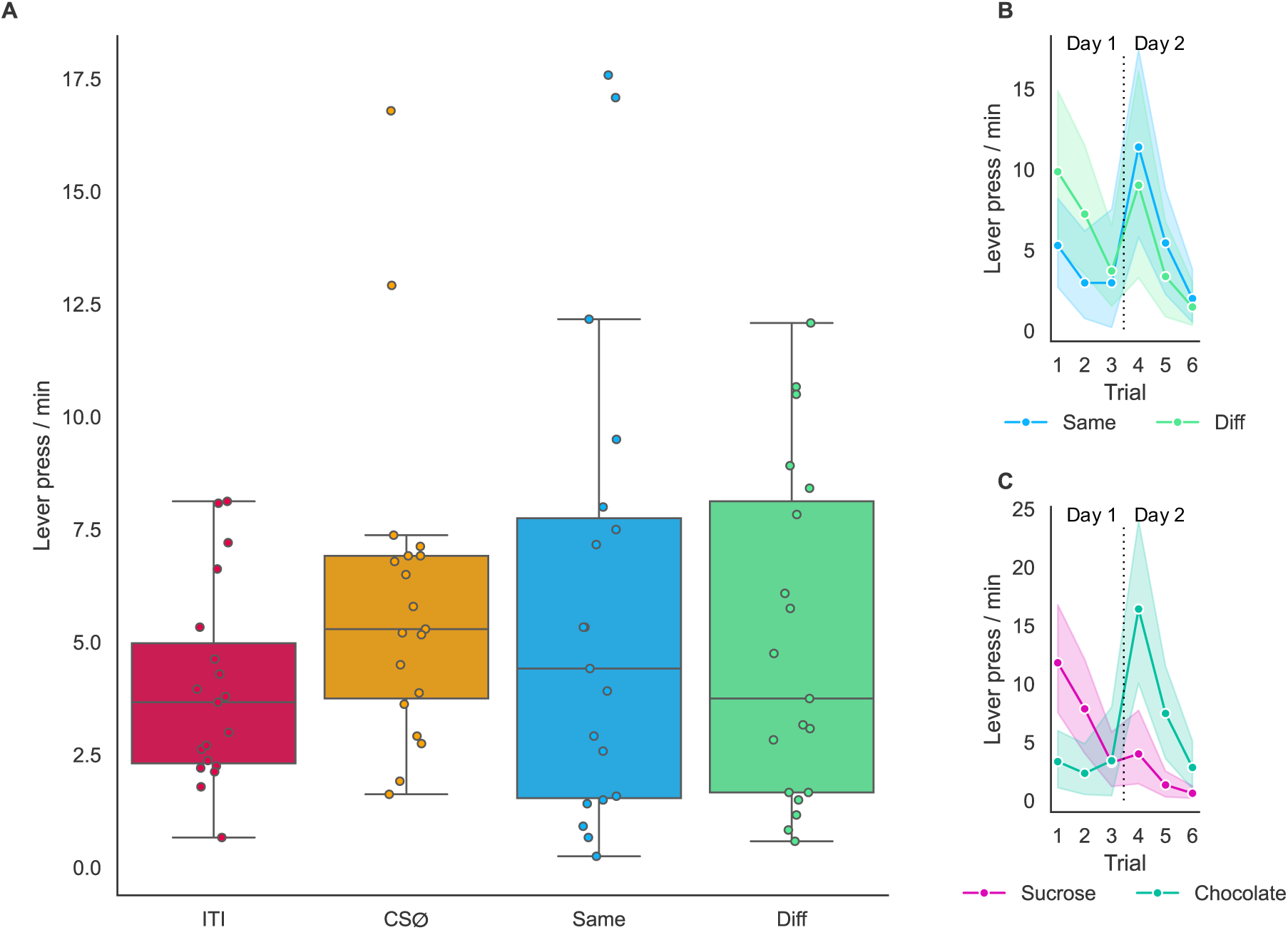
Instrumental extinction prior to the test reduces lever presses but female Lister-Hooded rats did not show transfer effects in a full transfer paradigm. Female Lister-hooded rats were trained on the full transfer paradigm PIT task, performed one instrumental extinction session and their performance was measured during the transfer phase. (A) The performance was classified based on the outcome, represented by intertrial interval (ITI), outcome-specific same, outcome-specific different (Diff) and general transfer effect. Lever presses total according to the PIT effect (B) and associated reward (C) across trials. Sample size n = 19. Data are shown as mean ± 95% C.I.

While there was no overall difference in responding for the CSs that had been associated with the two different rewards [**Fig. 7C**; Reward: *F*_(1, 18)_ = 1.18, *p* = 0.29, η_p_² = .062], responding for the CSs associated with sucrose decreased throughout the test, while responding for chocolate increased [Order: *F*_(1.47, 26.55)_ = 9.92, *p* = .002, η_p_^2^ = 0.36; Order x Reward: *F*_(1.97, 35.42)_ = 0.28, *p* = .75; Day x Reward: *F*_(1, 18)_ = 22.40, *p* < .001, η_p_^2^ = .55; Day x Order x Reward: *F*_(1.51, 27.24)_ = 8.13, *p* = .003, η_p_^2^ = .31].

### Sex differences in pavlovian and instrumental conditioning

To determine whether there were differences in pavlovian or instrumental learning in males and females, we compared the data from Experiments 1-3 (for training) and Experiments 2 and 3 (for transfer test; Experiment 1 was excluded from this analysis due to the lack of extinction prior to the test).

First, the pavlovian phase was analysed using only the stimuli that had been paired with rewards. All rats increased the number of nosepokes made during the rewarded cues (**Fig. 8A**) throughout training [Time: *F*_(17, 884)_ = 32.8, *p* < .001, η_p_^2^ = .39]. Although males and females started at a similar baseline, females rapidly increased the number of nosepokes made beyond the males [Time x Sex: *F*_(17, 884)_ = 2.62, *p* < .001, η_p_^2^ = .048; Sex: *F*_(1, 52)_ = 6.12, *p* = .017, η_p_^2^ = 0.11]. However, nosepoke duration was similar (**Fig. 8B**) between males and females, increasing in both sexes over the course of training [Time: *F*_(17, 884)_ = 16.9, *p* < .001, η_p_^2^ = .25; Sex: *F* < 1; Time x Sex: *F*_(17, 884)_ = 1.19, *p* = .27], suggesting that females entered the magazine more often, with no sex differences in duration.

**Figure 8.**
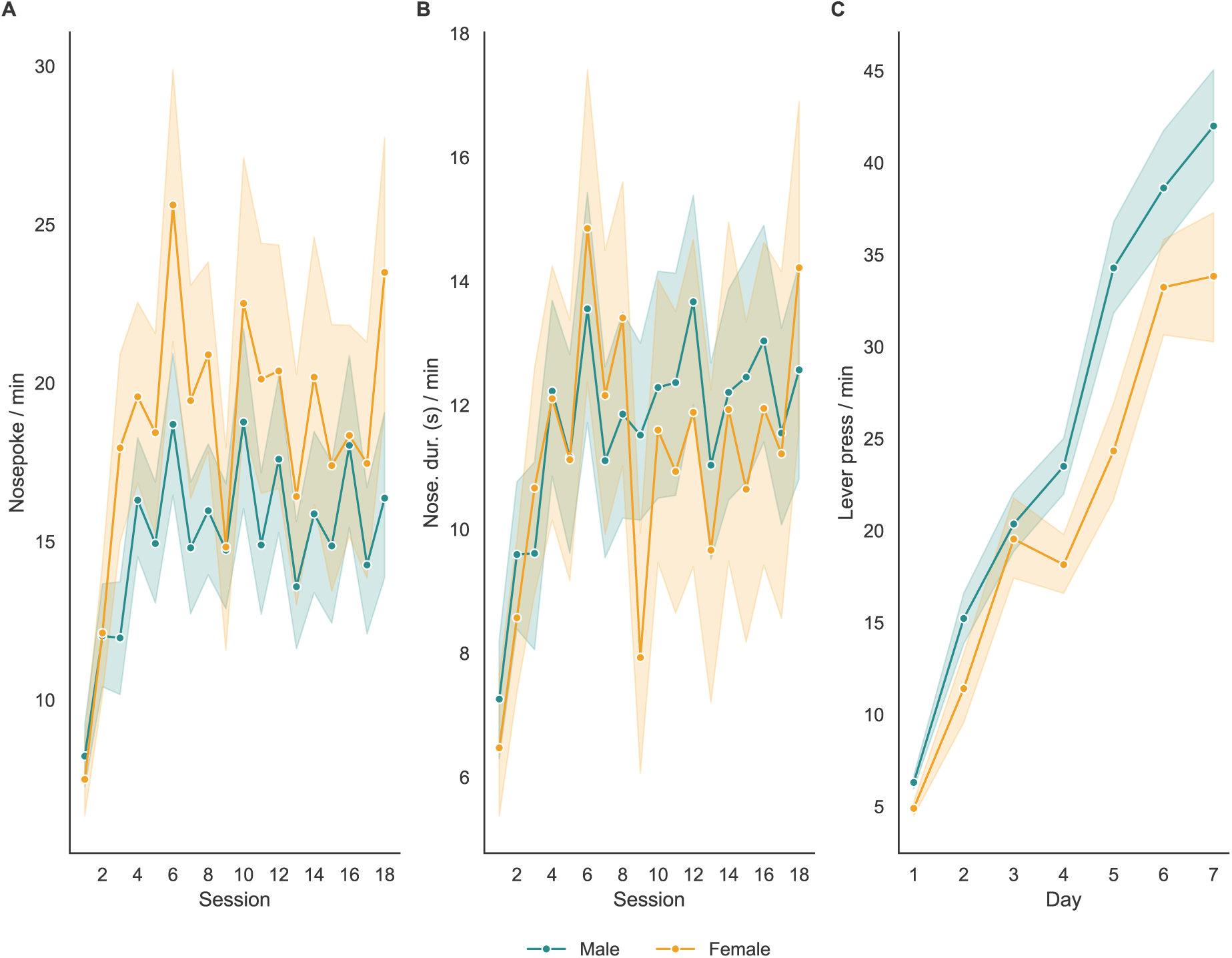
Female rodents perform more nosepokes than male, but with reduced lever pressing. All animals from experiments 1, 2 and 3 were grouped to compare performance for the pavlovian and instrumental phases. (A) Nosepoke entries and (B) duration for each session of the pavlovian phase. (C) The total amount of lever presses across seven days of the instrumental training. Sample size n = 55 (male = 36; female = 19). Data are shown as mean ± 95% C.I.

Although both males and females acquired instrumental responding [**Fig. 8C**; Time: *F*_(6, 318)_ = 288, *p* < .001, η_p_^2^ = .85], females showed lower levels of lever pressing than males, particularly during the second half of instrumental training [Sex: *F*_(1, 53)_ = 10.5, *p* = .002, η_p_^2^ = 0.17; Time x Sex: *F*_(6, 318)_ = 4.86, *p* = .0001, η_p_^2^ = .084]. Thus, females performed better on pavlovian conditioning, but worse than males on instrumental conditioning.

### Magazine entries during the transfer tests

One theory on PIT suggests that magazine visits comprise part of the chain of behavioural responses learned during pavlovian and instrumental training, forming CS-magazine and lever-magazine associations, respectively (van den Bos et al., 2004; Holmes et al., 2010). However, it is debated whether magazine approaches would positively correlate or compete with instrumental responses (Cartoni et al., 2016). As there were no transfer effects as measured by lever pressing in any of the three experiments, we conducted an exploratory analysis on the number of magazine entries during the transfer tests (**Fig. 9**). For all three experiments, there was greater nosepoking during presentation of the reward-associated CSs, as compared to the CSØ and ITI periods [Experiment 1 PIT: *F*_(1.76, 40.4)_ = 10.6, *p* < .001, η_p_^2^ = .32; Experiment 2 PIT: *F*_(2.49, 27.4)_ = 6.12, *p* = .002, η_p_^2^ = .36; Experiment 3 PIT: *F*_(2.20, 39.5)_ = 6.35, *p* = .001, η_p_^2^ = .26]. For all three experiments, responding during the Same and Different CSs was higher than responding during the ITI [Bonferroni-corrected pairwise comparisons, all *p*’s < .03]. Notably, despite the lack of instrumental expression, in all experiments the number of magazine entries was elevated for all outcome-specific PIT, with only the different PIT effect of Experiment 1 being positively correlated with lever pressing (Fig. S1).

**Figure 9.**
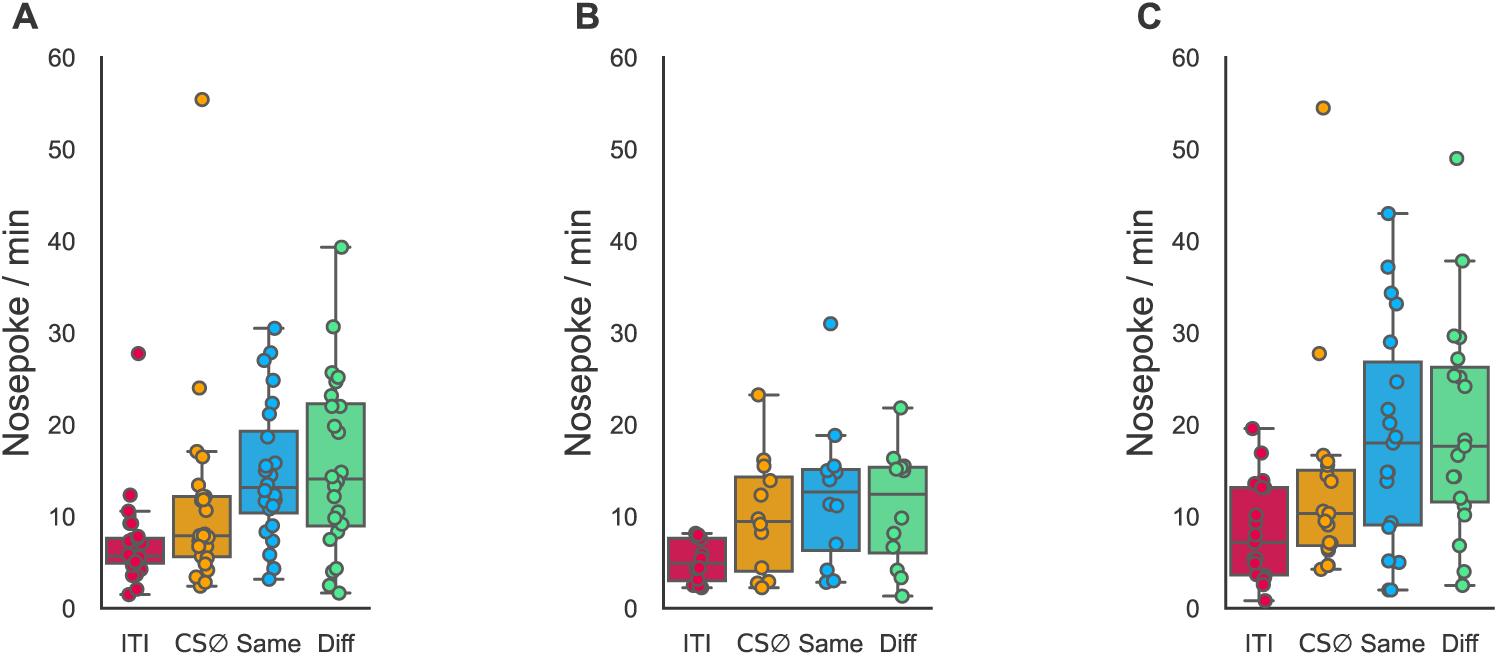
The number of nosepokes performed during the transfer phase is higher for both outcome-specific PIT in all experiments. (A) A one-way ANOVA pointed suggested a significant main effect for PIT effect (F(1.76, 40.42) = 10.61, p < 0.001). Follow up post-hoc comparisons using Bonferroni correction indicated a significant difference for all comparison with the intertrial interval (ITI vs CSØ: p = 0.03; ITI vs Same: p = 0.001; ITI vs Diff: p < 0.001). Sample size n = 24. (B) A one-way ANOVA pointed suggested a significant main effect for PIT effect (F(2.49, 27.44) = 6.12, p = 0.002). Follow up post-hoc comparisons using Bonferroni correction indicated that the rate of nosepokes during the intertrial interval was significantly different than both outcome-specific transfer (ITI vs Same: p = 0.03; ITI vs Diff: p = 0.03), with the active baseline CS being close to statistical significance (ITI vs CSØ: p = 0.05). Sample size n = 12. (C) A one-way ANOVA pointed suggested a significant main effect for PIT effect (F(2.20, 39.54) = 6.35, p = 0.001). Follow up post-hoc comparisons using Bonferroni correction indicated that the rate of nosepokes during the intertrial interval was significantly different than both outcome-specific transfer (ITI vs Same: p = 0.004; ITI vs Diff: p = 0.003). Sample size n = 19.

Comparing Experiments 2 and 3, while both males and females showed greater nosepoking during presentation of the reward-associated CSs [PIT: *F*_(3, 87)_ = 10.0, *p* < .0001, η_p_^2^ = .26], females continued to show higher levels of magazine entries, [Sex: *F*_(1, 29)_ = 5.27, *p* = 0.03, η_p_^2^ = .15], though in a similar response pattern to males during the CSs [Sex x PIT: *F* < 1].

## Discussion

Adaptive reward-seeking requires integration of predictive cues with learned actions to obtain desirable outcomes. PIT provides a critical framework for understanding how reward-paired cues motivates behaviour across species. The full transfer paradigm, in particular, has the advantage of being able to distinguish general and specific PIT effects (Corbit & Balleine, 2005; Corbit et al., 2007). However, to our knowledge, all animal designs lack a non-rewarded paired cue condition, making it difficult to evaluate whether PIT effects exist due to a facilitation of responding by a reward-paired cue or inhibition by the cue of incompatible actions. Combining the results from males and females, this work provides evidence that both successfully learnt the pavlovian conditioning on the modified version of the full transfer paradigm. Our data also suggests that both sexes learnt the instrumental conditioning showing consistency with previous work (Corbit et al., 2005, 2007). However, despite this robust learning, animals did not show transfer effects when our adapted full transfer paradigm was used, suggesting that the addition of a non-rewarded paired cue may alter the motivation underlying PIT.

In human PIT paradigms, neutral cues typically elicit baseline levels of responding comparable to intertrial intervals during both appetitive and aversive conditioning (Quail et al., 2017; Eder and Mitschke, 2025). Comparable results have been reported in rodents when a neutral cue is used sparingly (Collins et al., 2019). As outlined previously, the current study used an ‘empty’ reward as a reinforcer for CSØ, while others only used positive reward paired cues, which could be a sucrose or chocolate pellet. Under these conditions, we expected enhanced responding when the CS and instrumental action shared a common outcome and reduced responding when they did not. Contrary to this prediction, no outcome-specific PIT emerged under these experimental conditions, diverging from prior studies using a full transfer paradigm (Corbit et al., 2005, 2007, 2011; Derman and Ferrario, 2020; Panayi and Killcross, 2022). In fact, the present data suggest that non-reward predictive cues in the full transfer paradigm may act as a conditioned inhibitor, signalling the absence of reinforcement. Such inhibitory associations are known to suppress instrumental responses while leaving pavlovian responses intact (Kemp and Corbit, 2021). Consistent with this interpretation, animals maintained pavlovian responses such as magazine approaches during CSØ but failed to express instrumental transfer. Therefore, CSØ may have reduced the overall motivational value of the cues associated with the reward outcome.

Aspects of these data are also consistent with a framework provided by Balleine and Ostlund (2007). According to the two-process theory, during the instrumental phase, two associations are learned, an outcome-response and a response-outcome. Thus, during the test phase, the retrieval of the outcome by a pavlovian stimulus-outcome association would bias selection towards the instrumental response that is associated with that same outcome, resulting in an increase of instrumental response for outcome-specific same PIT. If CSØ acted as a conditioned inhibitor, it may have disrupted the retrieval of the outcome-response association, reducing lever pressing while preserving cue-elicited magazine approach behaviour. Prior work has shown that reward expectancy and cue–reward probability modulate transfer strength (Marshall et al., 2020, 2023), and that cues predicting low-value or more certain reward can suppress instrumental responding. Thus, the inclusion of a non-rewarded cue may have changed reward expectations, effectively reducing cue-guided motivational behaviour and masking transfer effects.

However, before accepting such conclusions, procedural and biological factors could also account for the absence of PIT in our modified paradigm. Methodological variables such as training duration and phase order are known to modulate PIT magnitude in both rats and humans (Holmes, et al., 2010; Colagiuri and Lovibond, 2015). Excessive instrumental training can elevate baseline responding and reduce transfer effects, whereas instrumental extinction prior to testing can reduce baseline rates, helping to restore PIT (Dickinson et al., 2000). Although instrumental extinction reduced responding in our study, it did not lead to transfer in either males or females, indicating that the lack of PIT was not due to elevated baseline performance. Instead, the elevated magazine entries observed during cue presentations positively correlated with lever pressing, suggest that magazine entry activity did not impose a limit on lever press performance. Thus, although animals had learned the associations between specific CSs, instrumental responses, and their corresponding rewards, this was not expressed by a modulation of instrumental behaviour.

Overtraining during Pavlovian conditioning may also interfere with transfer by inducing response competition (Holmes et al., 2010). More specifically, extensive pavlovian training (16 sessions) led to an apparent loss of PIT effects that were observed with more moderate pavlovian training (4 sessions). This effect may have occurred in our adapted design, due to the number of pavlovian conditioning sessions. To date, no systematic evaluation of these parameters has been conducted for the full transfer paradigm, underscoring the need for a comprehensive methodological analysis.

Biological variation may further shape PIT expression. The present study is, to our knowledge, the first to implement the full PIT transfer protocol in Lister-Hooded rats. The PIT effect using the full transfer paradigm was originally described in Long Evans rats (Corbit and Balleine, 2005). Likewise, Sprague-Dawley also showed transfer effects when the full transfer paradigm was used (Derman and Ferrario, 2020). In contrast, a recent study observed specific PIT effects in Wistar rats, but not general PIT (Panayi and Killcross, 2022). Within our experiments, our Lister-Hooded rats exhibited relatively high instrumental response rates during the instrumental phase in comparison to what was previously described in the literature (Corbit et al., 2007), suggesting strain-dependent motivational differences (Andrews et al., 1995; Waite et al., 2021). These data suggests that rodent strains may contribute to the expression of PIT effects.

Despite the possibility that stress-related factors could module PIT outcomes in both humans and rodents (Quail et al., 2017; Sommer et al., 2022; Morgado et al., 2012), this is unlikely to account for the observed lack of transfer. Previous studies showed that chronic stress can induce impairment of PIT outcomes in rodents (Morgado et al., 2012), while acute stress (Pielock et al., 2013) reduce basal instrumental responding during the test, both without interfering with pavlovian and instrumental learning. Our design included extensive habituation and handling procedures, as well as home-cage exposure to the rewards, to minimize novelty and handling stress (Greiner & Petrovich, 2020). Auditory cues were also presented at attenuated intensities relative to previous protocols (Corbit & Balleine, 2005) to reduce the probability of potential audiogenic seizures in the Lister Hooded strain used in our experiments (Commissaris et al., 2000). These measures make it improbable that arousal or stress effects masked the expression of PIT.

Sex is an important biological variable that can shape behavioural strategies with implications on the cue-driven motivation necessary during PIT. In the present study, sex differences also shaped performance. We observed differences in the propensity for all animals to express instrumental and pavlovian learning, though neither males nor females expressed PIT during the transfer test. This is partially consistent with a previous study that found no sex differences for pavlovian conditioning or in the transfer effects using a single-lever PIT procedure (Garceau et al., 2023). Moreover, this study did observe higher levels of instrumental responding in males compared to females, consistent with our observations. A study which evaluated the selectivity of PIT produced by CSs that signal alcohol also reported no sex differences between instrumental and pavlovian learning (Alarcón and Delamater, 2019). Similarly, in humans, no gender effect was observed on a pavlovian instrumental transfer test (Seabrooke et al., 2018). Overall, our findings are consistent with the evidence from other PIT paradigms, showing males pressing more than female rodents during the instrumental phase. However, during the pavlovian phase females performed more pellet dispenser entries in comparison with males, despite no significant difference on the amount of time spent in the pellet dispenser. If female rats, at least Lister-Hooded, better discriminate appetitive CS-outcome pairings on a more complex pavlovian task, perhaps less pavlovian training is required for the animal to express PIT. As noted previously, in a two lever PIT paradigm using both CS+ and CS-cues, initiating the pavlovian phase first and inducing overtraining can disrupt PIT expression (Holmes et al., 2010). This was accompanied with increased pellet dispenser visits relative to baseline.

In conclusion, our findings reveal that incorporating a non-rewarded cue as a control in the full transfer paradigm precludes PIT effects in both sexes, despite intact pavlovian and instrumental learning. We propose that CSØ acquired inhibitory properties that signalled non-reward and devalued motivational drive, preventing cue-induced facilitation of instrumental behaviour. By demonstrating that cue-driven motivation can be suppressed by the presence of a non-rewarded cue, our results uncover aspects of reward prediction and goal-directed behaviour. The inclusion of CSØ enhances the translational relevance of rodent PIT paradigms more consistent with human PIT methodologies (Quail et al., 2017), in which non-rewarded or neutral cues are routinely used to dissociate cue-induced facilitation from baseline inhibition. This new framework allows the assessment of how reward-predictive cues compete with inhibitory signals to guide behaviour, a mechanism relevant to both adaptive decision-making and its disruption in psychiatric disorders.

## Acknowledgments

FEA is supported by UK Research and Innovation’s Biological and Biotechnology Research Council grant BB/W001195/1. ALM is the Ferreras-Willetts Fellow in Neuroscience at Downing College, University of Cambridge.

---

**Figure S1.**
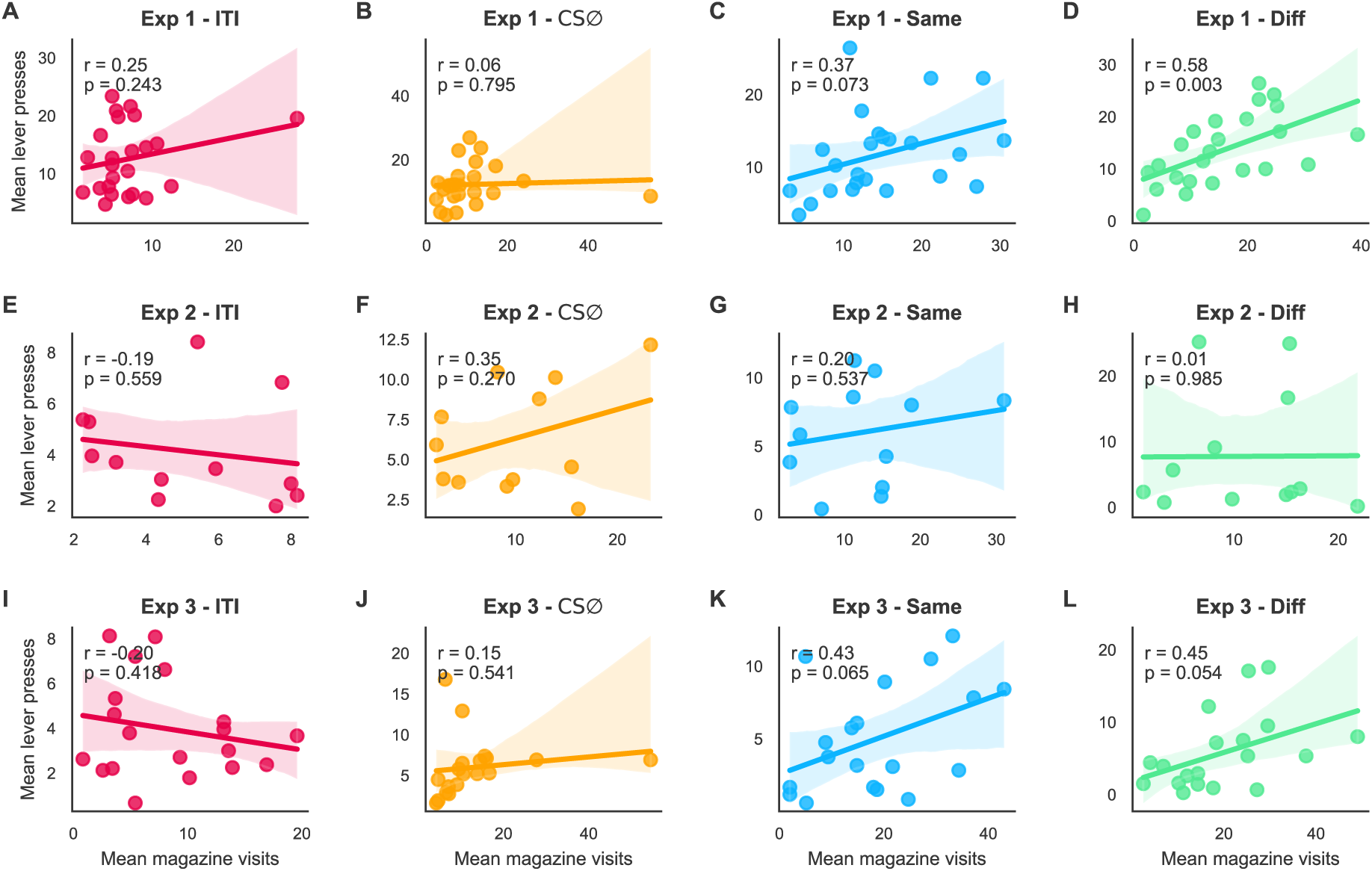
Correlation between average magazine visits and lever presses across PIT conditions and experiments. Each row represents one experiment (Experiments 1 to 3); each column corresponds to a PIT condition (ITI, CSØ, same, and different). Scatter points indicate mean values per rat, and regression lines (colored per condition) illustrate the correlation direction. Pearson’s correlation coefficients (*r*) and *p*-values are displayed within each panel.

